# Large extracellular vesicles subsets and contents discrimination: the potential of morpho mechanical approaches at single vesicle level

**DOI:** 10.1101/2025.03.29.646084

**Authors:** Geetika Raizada, Benjamin Brunel, Benoit Bragantini, Joan Guillouzouic, Eric Le Ferrec, Wilfrid Boireau, Eric Lesniewska, Céline Elie-Caille

## Abstract

Extracellular vesicles (EVs) are heterogenous lipid bound membranous structures released by different cells, showing a great potential to be used as biomarkers. They have also been explored for their role in the context of environmental toxicity. When endothelial cells are exposed to pollutants like Polycyclic Aromatic Hydrocarbons (PAH) - the most common being benzo[a]pyrene (B[a]P) – EVs released from those cells undergo surface and cargo modifications. Subpopulations of large EVs (lEVs) have shown to contain either damaged or intact mitochondria which is inexorably linked to oxidative stress conditions. In this paper, we studied B[a]P induced modifications in lEVs derived from endothelial cells, through morpho mechanical characterization with atomic force microscopy (AFM). Colocalizing AFM with fluorescence microscopy allowed us to differentiate between EVs containing mitochondria and those that did not. EVs containing mitochondria had a larger size (maximum diameter) when coming from treated cells (1.8 ± 0.89 µm) as compared to control cells (1.63 ± 0.76 µm). Moreover, their Young’s moduli were higher in the treated condition (3.09 ± 2.54 MPa in average) as compared to the control condition (1.25 ± 0.92 MPa in average). We also observed a heterogeneity within single vesicles, with most Young’s modulus values ranging from 0.1 up to 30 MPa for the treated condition and from 0.1 to 5 MPa for the control condition. Finally, applying linear discriminant analysis (LDA) and Random Forest (RF) algorithms on maximum diameter, height, and distribution of Young’s modulus values, we demonstrated the possibility to discriminate between EV subpopulations. Indeed, we successfully managed to a) distinguish EVs containing mitochondria from the “empty” ones, with an accuracy of 84% and b) discriminate whether these mitochondria-containing EVs originated from control or treated conditions, with an accuracy of 76%. These findings highlight the power of combining morpho-mechanical analysis and machine learning for identifying and discriminating EV subpopulations, no longer requiring any EVs fluorescence labelling.

## Introduction

Atomic Force Microscopy has been extensively used to study the morphology and mechanical properties of biological materials. It enables measurements under aqueous and physiological conditions with spatiotemporal resolution, thereby preserving the native state of biomolecules and features [1]. In AFM, a cantilever scans the sample surface while monitoring the interaction between the tip and the sample through force-distance curves. By exploiting these curves, we can extract viscoelastic and other mechanical properties of the samples, providing quantitative insights on the structural integrity of the biological samples [2], [3], [4].

Nanomechanical properties are often closely linked to the physiological and pathological state, as well as the content of biological samples. In recent years, AFM has been employed to study the extracellular vesicles (EVs) – a heterogenous population of lipid bilayer membranous structure that are released from different cells [5]. They play a crucial role in intercellular communication by carrying molecular cargo derived from their parent cells [6]. Their composition reflects the functional state of the cells of origin, making them valuable biomarkers for various diseases [7]. One of the major challenges in EV research lies in their pronounced heterogeneity—spanning a broad range of sizes, membrane and internal compositions, and mechanical properties [5]. AFM enables high-resolution, single-vesicle imaging and mechanical profiling, making it an ideal technique to resolve the inherent heterogeneity of EV sub-populations and distinguish co-isolated nanoparticles within complex biological samples. For example, Ridolfi et al. introduced a method of EV characterisation where they used the morphological data of EVs obtained by AFM to discriminate different subpopulations. They used contact angle of the individual EVs and size profiles to separate vesicles from the non-vesicular content in the sample [8], [9]. In other instance, blood derived EVs were characterized into subpopulations by PeakForce tapping® mode where the distribution of Young’s modulus within single vesicle was used to separate EVs subpopulations [10]. Leiva-Sabadini et al. studied the mechanical properties of EVs derived from honey where they measured Young’s moduli of small EVs (sEVs). They also used nanoindentation to study the antibacterial effect of sEVs on the membrane of Streptococcus mutans [11]. The study of metrology and nanomechanical properties has also been used for diagnostic applications. Sharma et al. showed the potential of using the morphology of saliva derived exosomes and the expression of CD63 as oral cancer biomarkers [12]. In another recent study, AFM was used to obtain nanomechanical signatures of the EVs coming from bone marrow and blood of hematologic cancer patients. They showed a clear difference between multiple myeloma patients, lymphoma patients and healthy donors [13]. In our previous work, we combined AFM with surface plasmon resonance imaging (SPRi) to investigate the morphological characteristics of EVs selectively captured by antibodies immobilized on an SPR-based biochip. This hybrid approach enabled us to distinguish EV subpopulations according to the targeted ligands and cell culture conditions. Notably, statistical analysis of AFM-derived morphological parameters revealed distinct EV groups characterized by variations in size and protein expression profiles, emphasizing the importance of quantitative approaches for capturing biologically meaningful heterogeneity [14], [15]. However, when extending such analysis to nanomechanical characterization, the complexity increases substantially, as each AFM map may contain thousands of force– distance curves. Given the heterogeneity of EV subpopulations, manual or semi-quantitative processing of these datasets can lead to significant information loss and limits the reliability of the mechanical interpretation. To investigate how such heterogeneity manifests under biologically relevant stress conditions, we focused on extracellular vesicles released by endothelial cells exposed to cytotoxic stimuli.

In this study, we investigated EVs coming from Human microvascular endothelial cell line (HMEC-1) which are exposed to benzo[a]pyrene (B[a]P). B[a]P is one of the polycyclic aromatic hydrocarbons (PAH) that can be found in diet, smoke, water, etc. When cells are exposed to it, it is known to generate oxidative stress [16] resulting from an influx of reactive oxygen species (ROS) [17] which then causes processes such as lipid peroxidation, DNA damage, etc [18]. Exposure to these toxicants could lead to changes in EV composition, cargo and proteins [19]. There have been several studies which explored EVs as biomarkers for cytotoxicity and their role in promoting oxidative stress [20], [21]. The rise in the production of ROS under stress condition could lead to mitochondrial damage as it is intricately linked with the metabolism of ROS [22]. Damaged mitochondria can be removed from the cells by different cellular processes such as mitophagy [23]. It has been also shown previously that mesenchymal stem cells derived EVs transferred intact mitochondria inside microvesicles under intracellular oxidative stress [24].

Building on these findings, our experimental model was designed to investigate the effect of cytotoxic stress on the morphological and mechanical properties of large extracellular vesicles (lEVs), with particular focus on the presence of mitochondria within individual vesicles. Fluorescence microscopy was employed to identify lEVs containing mitochondria, providing a reference framework for targeted AFM-based morpho-mechanical analysis. By integrating AFM-derived morphological and nanomechanical parameters with statistical and machine-learning approaches, we aimed to discriminate between EV subpopulations containing or lacking mitochondria and to assess differences arising from control and B[a]P-treated conditions. This integrated strategy highlights the potential of combining AFM with data-driven analysis to identify and classify EV subtypes objectively, offering the possibility of eliminating the need for fluorescent labelling in the future, and to deepen our understanding of EV heterogeneity under stress-induced cellular conditions.

## Method

### 1. Cell culture and EV isolation

In this study, we employed the HMEC-1 cell line, a human microvascular endothelial cell line, sourced from the Center for Disease Control and Prevention, Atlanta, USA. The cells were cultured in endothelial basal medium MCDB131 (US Biological Life Sciences, Ref E3000-01G) containing 10% decomplemented (56°C, 30 min) fetal bovine serum (FBS, Dutscher, Ref 500105A1A), L-glutamine (Gibco, Ref 25030, 10 mM final), penicillin/streptomycin (Gibco, Ref 15140-122, 100 unit/mL final), gentamycin (Gibco, Ref 15750-037, 500 µg/mL final), epithelial growth factor (EGF, Sigma, Ref E9644, 10 ng/mL final) and Hydrocortisone (Up John, Ref 3400932141159, 1 µg/mL final) in 151.9 cm^2^ petri dishes (Corning, Ref 353025). The pH of the medium was adjusted at 7.6 using sodium bicarbonate (Gibco, Ref 25080-060). The medium was replaced every two to three days. Cells underwent weekly passage via trypsinization (trypsin EDTA 0.05%, Gibco, Ref 25300-054). When 90% confluence was achieved, we cultured cells overnight in FBS free medium. Later cells were treated with B[a]P 100 mM (Benzo[a]pyrene, Sigma, Ref B1760) dissolved in DMSO (DMSO, Sigma, Ref D8418), the final concentration of DMSO did not exceed 0.0005% (v/v) (treated condition) and control cell culture were exposed to the same amount of DMSO (control condition).

The method of large EV isolation was previously described by Le Goff et al. [25] which followed the international guidelines for extracellular vesicles isolation [5]. The FBS free medium was centrifuged at 3650 x g for 10 mins at 4°C, this step was done to remove cells and cell debris. Mito Tracker Green FM (MTG) (Thermo Fisher Scientific, USA) at a concentration of 100 nM was used to label the mitochondria inside the EVs. It was added to the preheated clarified medium (37°C) and was then incubated at 37 °C away in the dark for 45 mins. After the incubation, the clarified medium was ultracentrifuged at 10,000 g for 30 mins at 4°C (Optima L-90 K ultracentrifuge and Sw 28.1 Ti rotor or XE-90 ultracentrifuge and Sw 32 Ti rotor, Beckman Coulter, USA). The obtained pellet was washed by centrifuging it in PBS 1X buffer (Gibco, 14190-094) for 30 mins at 4°C. Finally, the pellet was resuspended in 50 µl PBS 1X buffer to be stored at −20°C.

Nanoparticle tracking analysis (NanoSight NS300, Malvern Instruments, UK) was performed to obtain the concentration of the EVs with the protocol of 5 videos of 60 s (Results: Figure S1). EVs from the treated condition will be referred to as t-EVs-M (treated EVs containing mitochondria) and t-EVs (treated EVs without mitochondria). For the control condition, they will be referred to as c-EVs-M (control EVs containing mitochondria) and c-EVs (control EVs without mitochondria).

### 2. Substrate preparation and sample deposition

All morpho mechanical measurements were performed on freshly cleaved mica. Since mica inherently have negative charges, we introduced mica sheets with divalent positive ions right after cleaving. The mica sheets (Agar Scientific, Ref: AGG250) were first cleaved with a piece of Scotch tape, then we quickly added 50 µl HEPES 10mM (Sigma-Aldrich, H3375-100G) + CaCl_2_ 5 mM (Sigma-Aldrich, 223506-500G) buffer solution onto the cleaved surface. After incubating the sheets for approximately 10 mins at room temperature, 5 µl of the sample solution containing EVs was added to the 25 µl buffer droplet and mixed gently by aspirating the solution at three different places. This step allowed us to distribute the EVs on the mica surface as uniformly as possible and to avoid forming exceptionally large aggregates. Then we left the mica sheet with EV sample to incubate for another 10 – 15 mins. Afterwards, we gently removed 25 µl from the buffer droplet to eliminate any floating objects that were not adsorbed on the mica surface. We then added 25 µl of buffer to further imaging under AFM.

### 3. Optical calibration between fluorescence and atomic force microscopy

The prepared mica sheet was observed under fluorescence microscope using 40x objective at an excitation wavelength of 480 nm to observe the EVs containing mitochondria. The laser was aligned on the top of the AFM tip, and the optimal SUM signal was obtained. After the tip approached the sample surface, it was retracted by up to 20 µm to keep it within the field of view during optical calibration. AFM apparatus (Bruker Nanowizard®4 Bioscience system) allowed the optical calibration by using the feature DirectOverlay which allowed us to determine whether the scanned area contained EVs with mitochondria or not. The calibration was done by acquiring the optical image of AFM tip at 25 or more different positions. The coordinates of tip at each position were automatically determined by AFM closed loop scanning system. These coordinates helped in accurately overlaying AFM image on the top of the fluorescence image. After the calibration step, we took a snapshot image to be used as a guide for scanning. After acquiring one or two AFM images, the corresponding shifted optical image was used to fine-tune the alignment between the optical and AFM channels. This calibration enabled precise identification of mitochondria-containing EVs (Fig 1).

**Figure 1.**
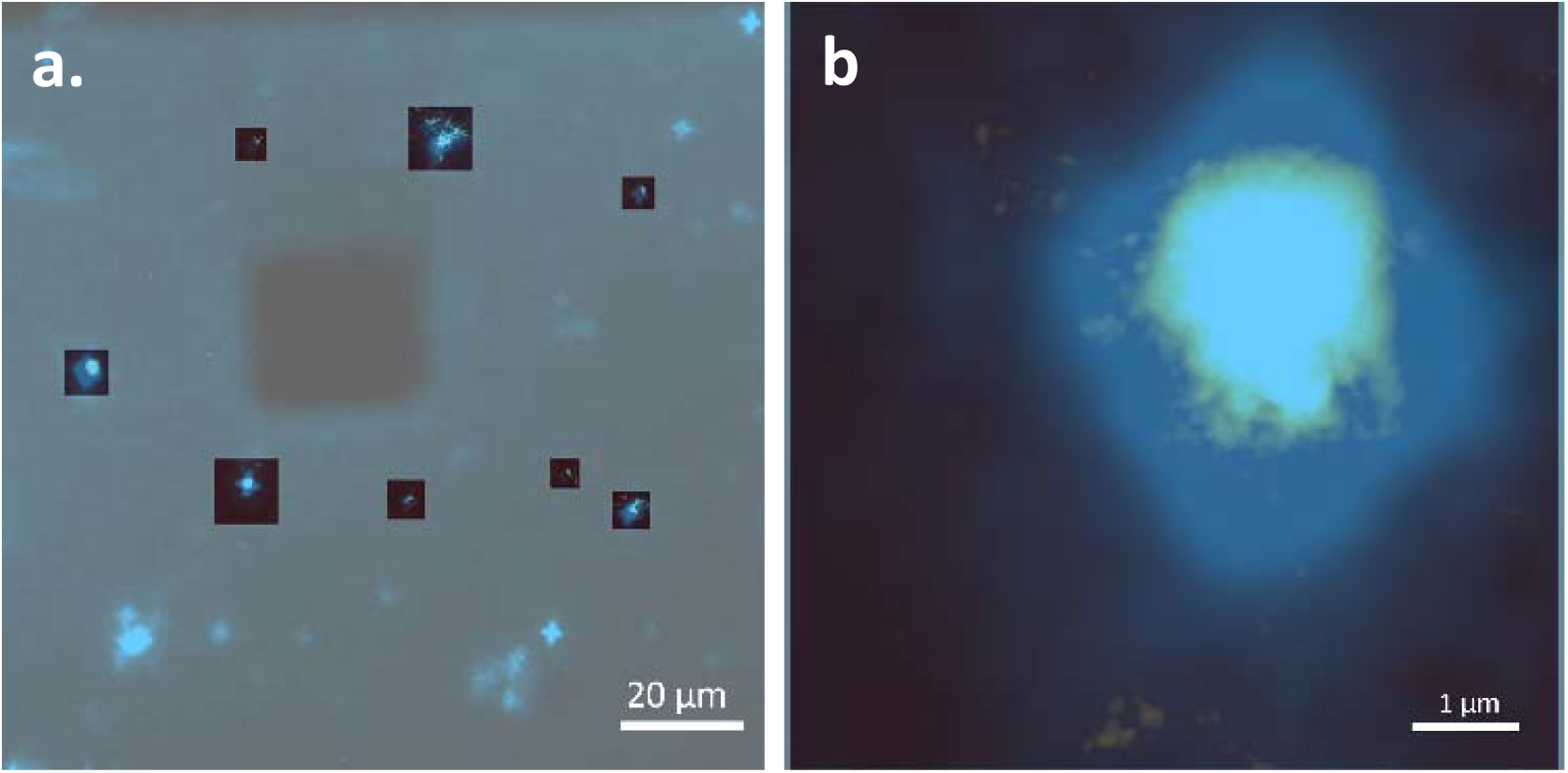
EVs derived from HMEC cells, adsorbed on mica, imaged by AFM coupled with fluorescence microscope. AFM images overlaid on top of fluorescence image obtained using 40x objective. a) Fluorescence image with overlay several AFM images; b) Focused AFM image of EV containing mitochondria with the fluorescent spot in the background.

### 4. Quantitative Imaging (QI) measurements

All quantitative imaging (QI) measurements were performed using QP-BIO-AC cantilevers (Nanosensors, Switzerland). These cantilevers, made of silicon or silicon nitride, are well suited for vesicle imaging and indentation experiments due to their low spring constant (0.06–0.3 N/m) and tip radius of approximately 10 nm. The pyramidal tip geometry with a rounded apex provides a good balance between resolution and gentle interaction with soft biological samples [26]. All imaging was carried out in liquid to preserve the physiological conditions of the EVs during QI-mode operation. In this mode, the force-distance curves (Fig 2) at each pixel are recorded which would enable us to calculate Young’s modulus of the objects scanned. The imaging was performed with a resolution of 256 x 256 pixels at the constant tip velocity. Low setpoint values were used to ensure that we do not rupture the EV membrane while keeping the indentation at about 1-2 %. The setpoint value was kept between 0.3 – 0.5 nN except for the cases where high setpoint was required. The area of scanning was dependent on the region selected on the fluorescence image, usually between 2 x 2 to 5 x 5 µm^2^ but not exceeding 10 x 10 µm^2^.

**Figure 2.**
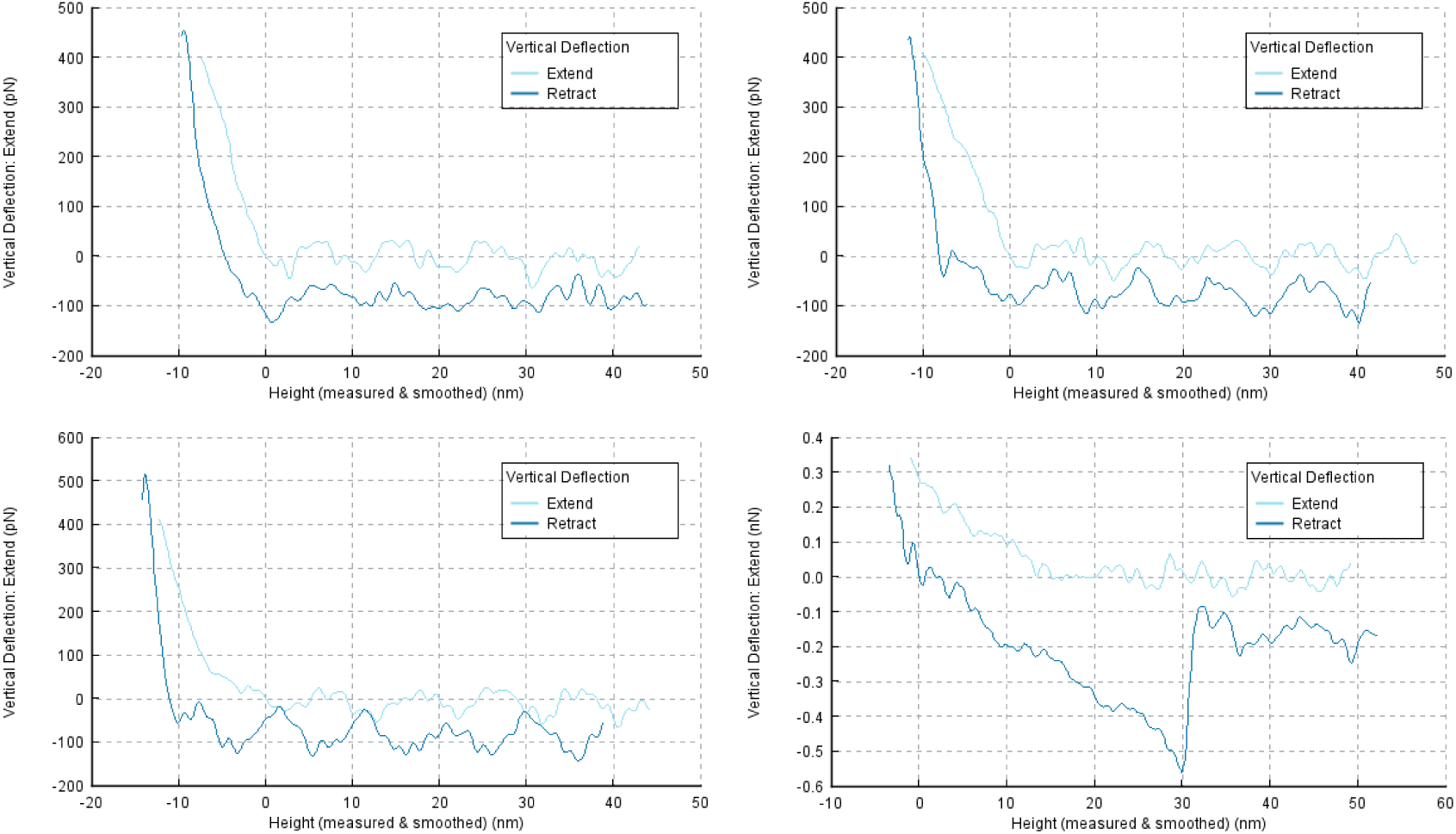
Examples of force-distance curves registered on several large EVs, obtained using QI mode in liquid. Approach (light blue) and retract (dark blue) curves. Tip used: QP-BIO-AC; Spring constant: 0.06 - 0.3 N/m; Tip radius: 10 nm.

### 5. Image treatment, data extraction and indentation model

All images and QI data were treated using Mountain’s SPIP 9 software. To obtain the size profile of the EVs, particle analysis was performed using a height threshold of 8.5 nm and a minimum area criterion of 0.2 µm^2^, which helped exclude free proteins and other elements that might have been adsorbed on the substrate from the sample solution, ensuring they were not considered during analysis [14], [27]. We first extracted force-distance curves at multiple points on EVs covering a broad surface area as shown in Fig 3, while avoiding all the points that are at the edge of the EV. Nanomechanical measurements were thus conducted at several positions rather than solely at the vesicle apex. This strategy was adopted to increase the likelihood of detecting mitochondria—whose dimensions and abundance can vary considerably—and to capture local variations in membrane mechanics. Measuring at multiple points allowed us to account for potential intra-vesicular and membranous heterogeneity, which could otherwise obscure distinctions between EV subpopulations.

**Figure 3.**
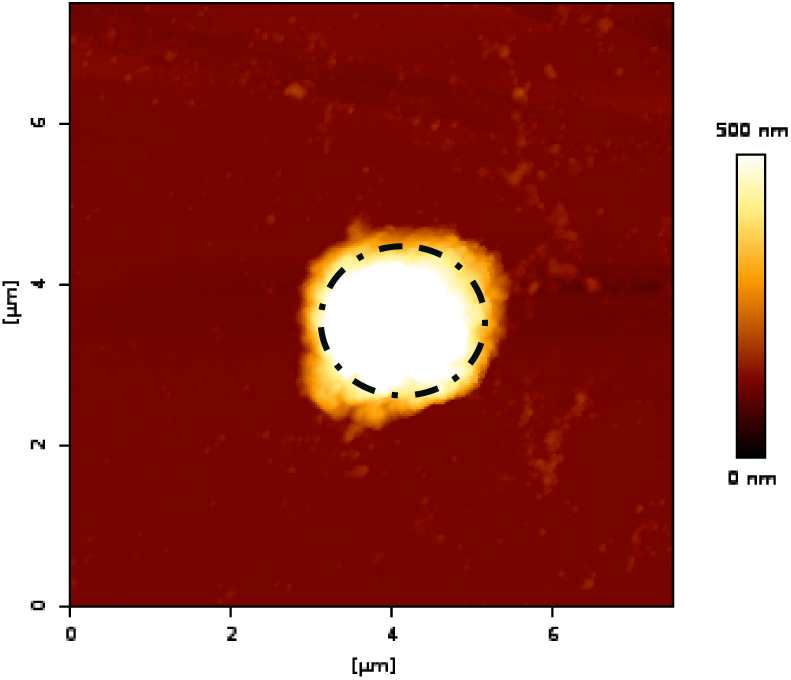
AFM (height channel) image showing one large EV; the region within the dotted line shows where we extracted the force-distance curves from several points (avoiding the edge of the vesicles).

The force–distance curves were subsequently smoothed using a Savitzky–Golay filter, followed by baseline correction through subtraction of a first-degree polynomial. The obtained curves were then calibrated by applying the values of photodiode sensitivity (the ratio between the tip deflection and the change in voltage output) and cantilever’s spring constant (k); this step allowed the measurement of the forces between the tip and the surface. To calculate the elastic modulus (Young’s modulus), we applied Derjaguin–Muller– Toropov (DMT) model, appropriate for a spherical AFM tip [28] using Poisson’s ratio as 0.3 for the analysis.

### 6. Statistical Analysis

Linear discriminant analysis (LDA) was performed on the Young’s modulus distribution (proportion of values < 1 MPa, between 1 and 5 MPa, and > 5 MPa), maximum height, and the maximum diameter to see the discrimination between different EV subpopulations measured. Analysis of variance (ANOVA) was performed on the LD1 and LD2 scores obtained, it was done to verify if there was a significant difference present between different populations or not. We applied the supervised Random Forest (RF) algorithm to distinguish EV subpopulations based on morpho-mechanical data, using the same assumptions as in the LDA analysis. The model’s performance was evaluated using leave-one-out cross-validation (LOOCV), ensuring unbiased discrimination and robust evaluation on this dataset. We used the ‘scikit-learn’ and ‘SciPy’ python libraries for LDA, Random Forest and ANOVA.

### 7. Cryo-EM characterization

4 µL of EV samples were applied on previously glow discharged (Pelco EasiGlow) C-Flat gold grids 1.2/1.3 300 mesh (Electron Microscopy Sciences) at concentrations of 10^10 particles/mL. Grids were then vitrified using a Vitrobot (ThermoFischer Scientific) in liquid ethane. The vitrification was performed at 4°C and 100% humidity, with a blotting time of 0.5 second. Cryo-EM images were acquired on a 200 kV Glacios 2 cryo-electron microscope from ThermoFischer Scientific equipped with a Ceta-D camera, using EPU software.

## Results and discussion

### 1. Morphological study of different conditions

The nanoscopic resolution offered by AFM has enabled us to closely examine the morphology of EVs derived from two different culture conditions i.e., control and treated. In our work, we focused primarily on the analysis of large EVs, those with diameters exceeding 200 nm. It must be noted that the different substrates and sample preparation techniques could influence the shapes of the EVs. The EVs isolation at 10,000 g centrifugation yielded a relatively heterogeneous population enriched in lEVs, which was intentionally preserved to maintain the natural diversity of vesicles prior to downstream morpho-mechanical and statistical sorting analyses. In our previous study, immunocapture experiments using antibodies against CD81, CD63, CD9, phosphatidylserine (Annexin-V) confirmed the presence of classical EV markers in vesicles derived from the HMEC-1 cell line. AFM imaging of these immunocaptured EVs revealed a heterogeneous size distribution for both control and treated conditions, consistent with the results obtained here [15].

For the present experiments, we utilized mica sheets treated with bivalent positive ions, which provide a remarkably flat surface having the average roughness Ra of 914.4 pm. The positive charge of the substrate surface facilitated the adsorption of the inherently negatively charged EVs without any additional selective process [29]. In most AFM images, we predominantly observed EVs with spherical geometries (as shown in figure 4a, b, c, and d).

**Figure 4.**
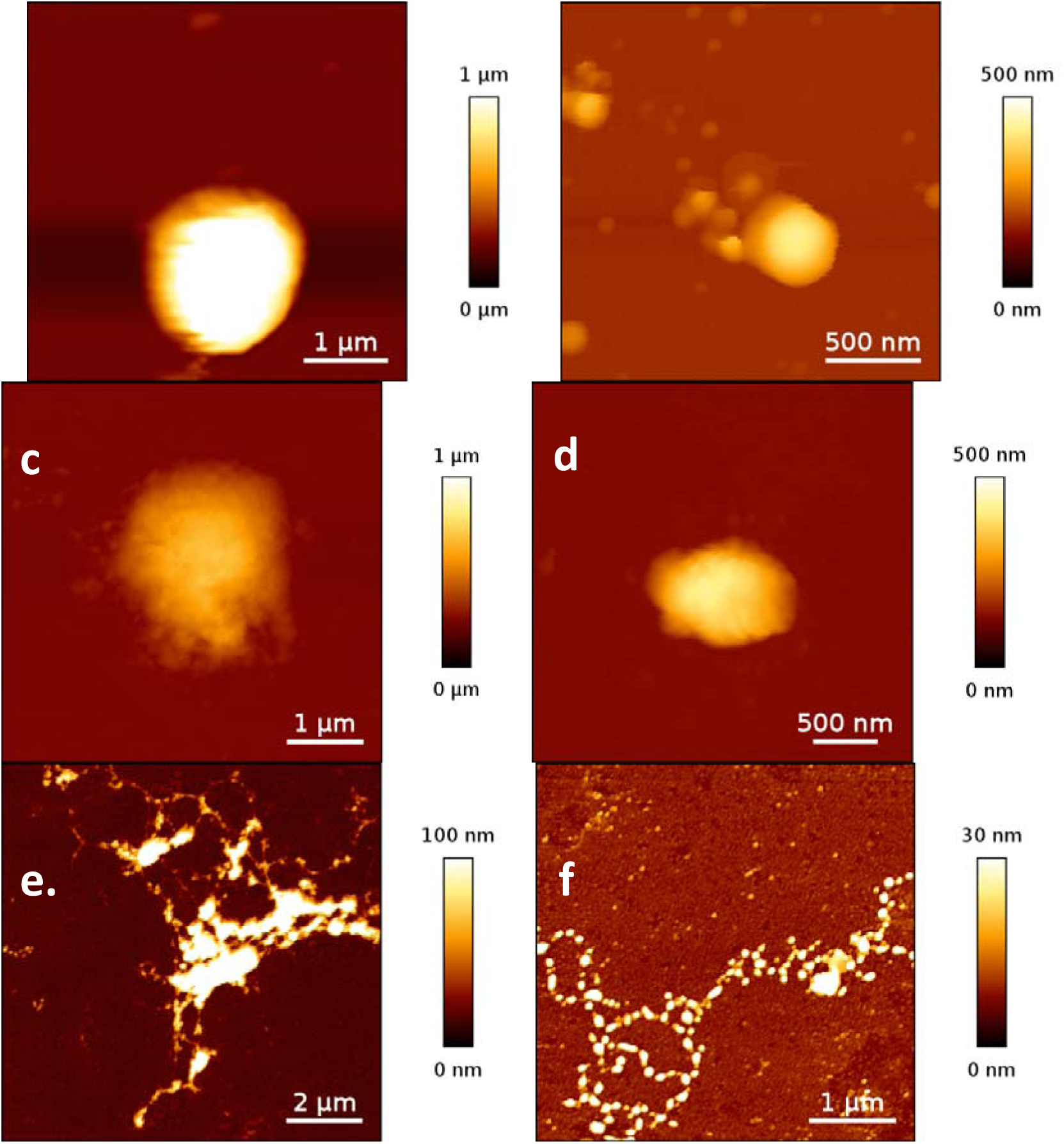
Morphological structure observed by AFM. a and b: lEV from control condition, c and d: lEV from treated condition, e and f: potential featurese that could be migrasomes and rectractosomes from treated condition respectively.

Notably, under the treated condition, we detected small vesicle-like structures clustering around and connected through fibres with lEVs (figure 4e). The lEVs in this structure ranged from 0.3 to 1.3 µm in size and were observed to contain mitochondria, a finding confirmed by fluorescence imaging (Figure S2). In another instance as shown in figure 4f, we observed a bead-like structure. These bead-like formations displayed heights ranging from 20 nm to approximately 50 nm, with profile sections closely resembling those of small EVs. These observations led us to hypothesize that these structures could correspond to migrasomes containing mitochondria [30], [31], with the smaller vesicle clusters possibly representing retractosomes [32].

Using AFM, we were able to conduct detailed size profiling of the EVs under study. Employing Mountain’s SPIP software for the analysis, we assessed EV size based on their maximum diameter and maximum height, as depicted in figure 5a and b. Figure 5a illustrates the distribution of EV sizes in terms of their maximum diameter, highlighting the presence of distinct EV subpopulations in both experimental conditions. The diameters of the EVs ranged significantly, reaching up to 5 µm. EVs devoid of mitochondria tended to be smaller on average than those that contain mitochondria. This observation was supported by NTA measurements (Figure S1) and further confirmed by cryo-EM analysis of the same samples (Figure S3).

**Figure 5.**
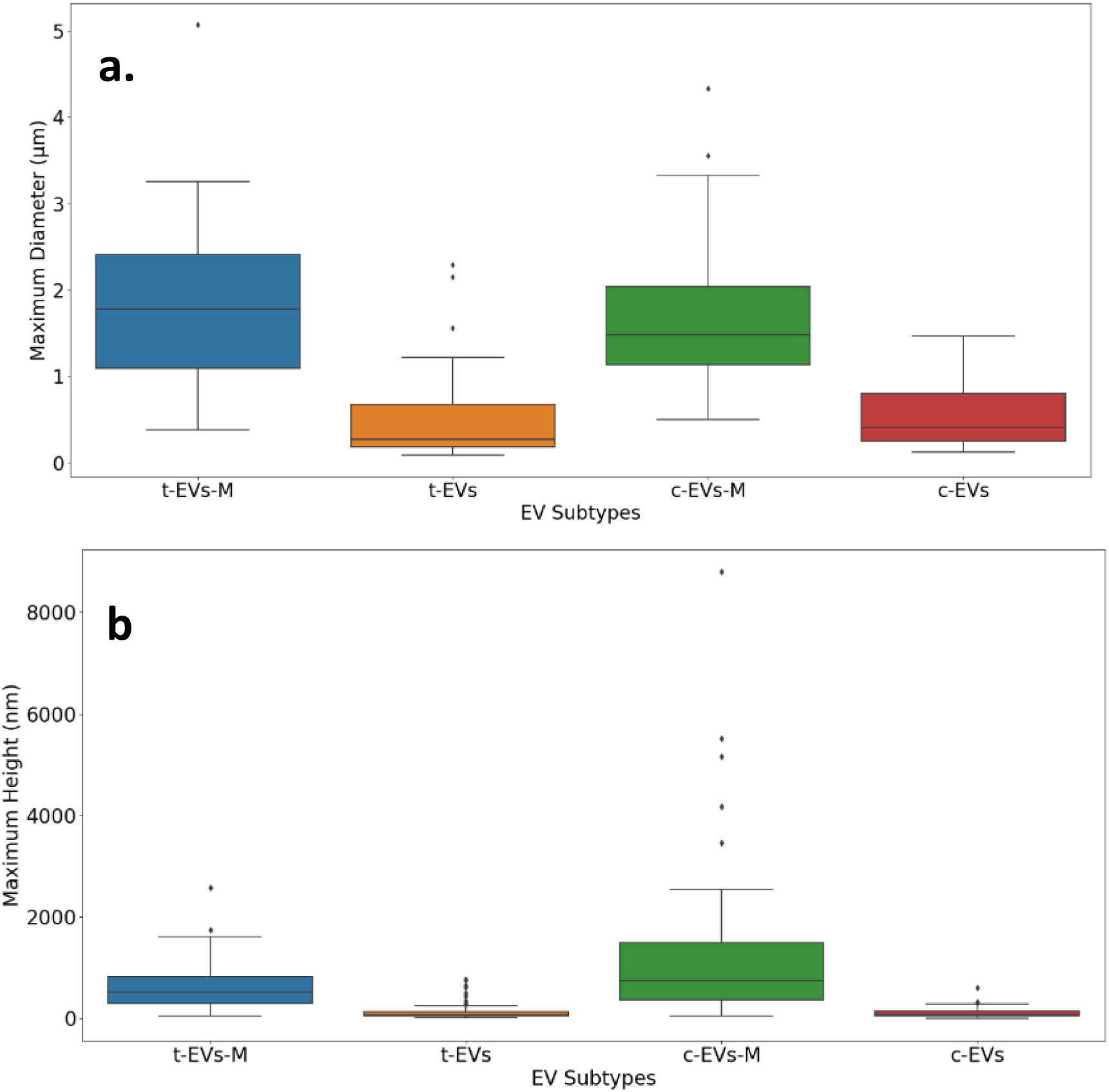
Size distribution of EV subtypes. a. box plot of maximum diameter and b. box plot of maximum height. For the size distribution, we measured 51 t-EVs-M, 69 t-EVs, 59 c-EVs-M and 53 c-EVs.

The mean maximum diameter calculated for c-EVs was 0.55 ± 0.37 µm and 0.47 ± 0.45 µm for t-EVs. From the box plot in figure 5b, it is evident that t-EVs-M did not exhibit significantly different maximum diameter from the c-EVs-M. Specifically, the average maximum diameter for the t-EVs-M is 1.8 ± 0.89 µm, whereas the c-EVs-M have an average maximum diameter of 1.63 ± 0.76 µm. After performing ANOVA on the maximum diameter of the lEVs with mitochondria (t-EVs-M and c-EVs-M) from both conditions, we obtained the p-value of 0.099, which indicates the differences between the size of lEVs from the two conditions was not statistically significant. This observation aligns with the study done by Le Goff et al., indeed they also did not observe significant change in the size of lEVs coming from B[a]P treated condition [25]. Additionally, we extracted the height of the EV subtypes (figure 5b) where we observed c-EVs-M were taller than t-EVs-M, with the means being 124_a_1 ± 1510 nm and 649 ± 491 nm respective_b_ly. Since we used positively charged surface and EVs are negatively charged, the observed height differences might be attributed to structural modification caused by B[a]P treatment. This data highlights the differences in size distributions among various EV subtypes under different conditions.

### 2. Nanomechanical discrimination

The use of QI mode in AFM allowed us to investigate the mechanical properties of EVs. We utilized the geometry of the AFM tips in the indentation model (DMT) to generate Young’s modulus maps as depicted in figure 6. The Young’s modulus maps provided us with the preliminary understanding of the mechanical properties of the various EV subpopulations present in both samples.

**Figure 6.**
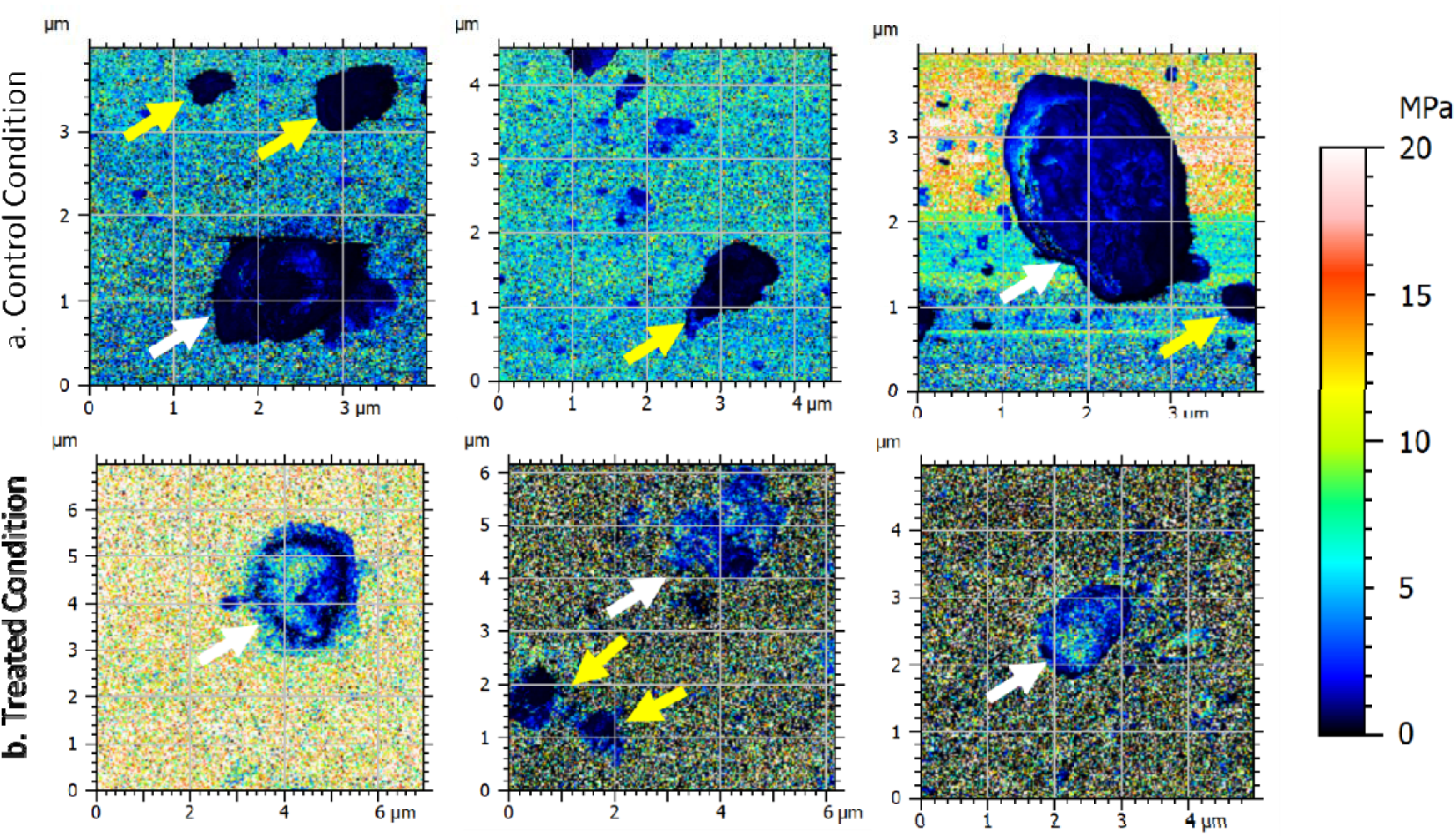
Young’s modulus maps generated after applying DMT indentation model, on lEVs from control and treated conditions. a. from control condition, b. from treated. The white arrows indicate EVs containing mitochondria and yellow arrows indicate EVs without mitochondria.

Based on young’s modulus maps, we can observe that c-EVs-M (figure 6a) tended to be softer than the ones from the t-EVs-M (figure 6b). EVs without mitochondria from both conditions (figure 6 shown with yellow arrows) tended to exhibit comparable elasticity. The difference in the elasticity of EVs from different conditions could be due to the modification of lEVs surface and cargo. In the oxidative stress conditions, EVs have shown to have both beneficial and deleterious effect of ROS metabolism [33]. In both cases, EVs go through a modification of their content either by carrying antioxidant machinery or due to processes such as lipid peroxidation which can induce changes in the membrane fluidity [34].

Following a qualitative discrimination of the EVs, we proceeded to extract the force-distance curves from individual EVs. This step was essential to establish the distribution profile of Young’s modulus values across single EVs. By analysing these curves, we aimed to gain a deeper understanding of the mechanical properties and variability of the EVs at a single-vesicle level. For each individual vesicles, we extracted several values of Young’s modulus, which were categorized into three distinct ranges: < 1 MPa, 1-5 MPa, and > 5 MPa (Note that the sum of the 3 proportions is always 100 % which explains why all the dots in fig 7 are inside a plane). This categorization allowed us to visualize systematically the distribution of Young’s modulus of individual EVs. To further cluster the EV subpopulations according to their mechanical properties, we used the 3D plot, which effectively illustrated the distribution and frequency of the Young’s modulus values across the different categories (Figure 7a). Figure 7a highlights the distinct elasticity profiles among various EV subpopulations. t-EVs-M (represented by blue dots) exhibited greater young modulus values as it has the higher proportion of values exceeding 5 MPa. In contrast, c-EVs-M showed comparatively lower young modulus values, as it contained less proportion of values surpassing 5 MPa. Additionally, the Young’s modulus distribution profile of EVs without mitochondria from both conditions (t-EVs and c-EVs) demonstrated the tendency to have Young’s modulus values smaller than 5 MPa (orange dots for treated condition and red for the control condition). Through these observations, the distinct clusters of EV subpopulations, differentiated by their mechanical properties, can be visualized in the 3D plot.

**Figure 7.**
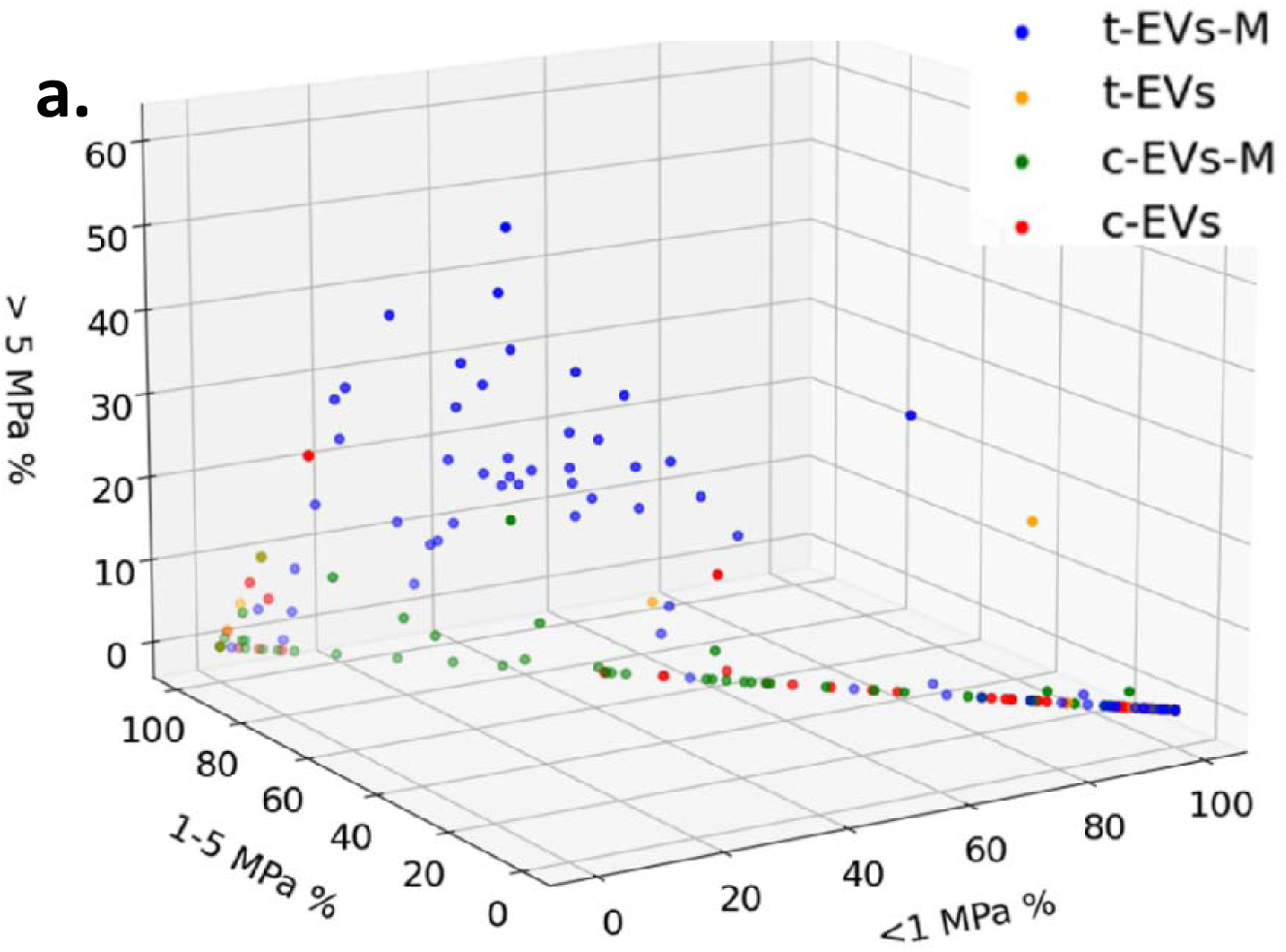
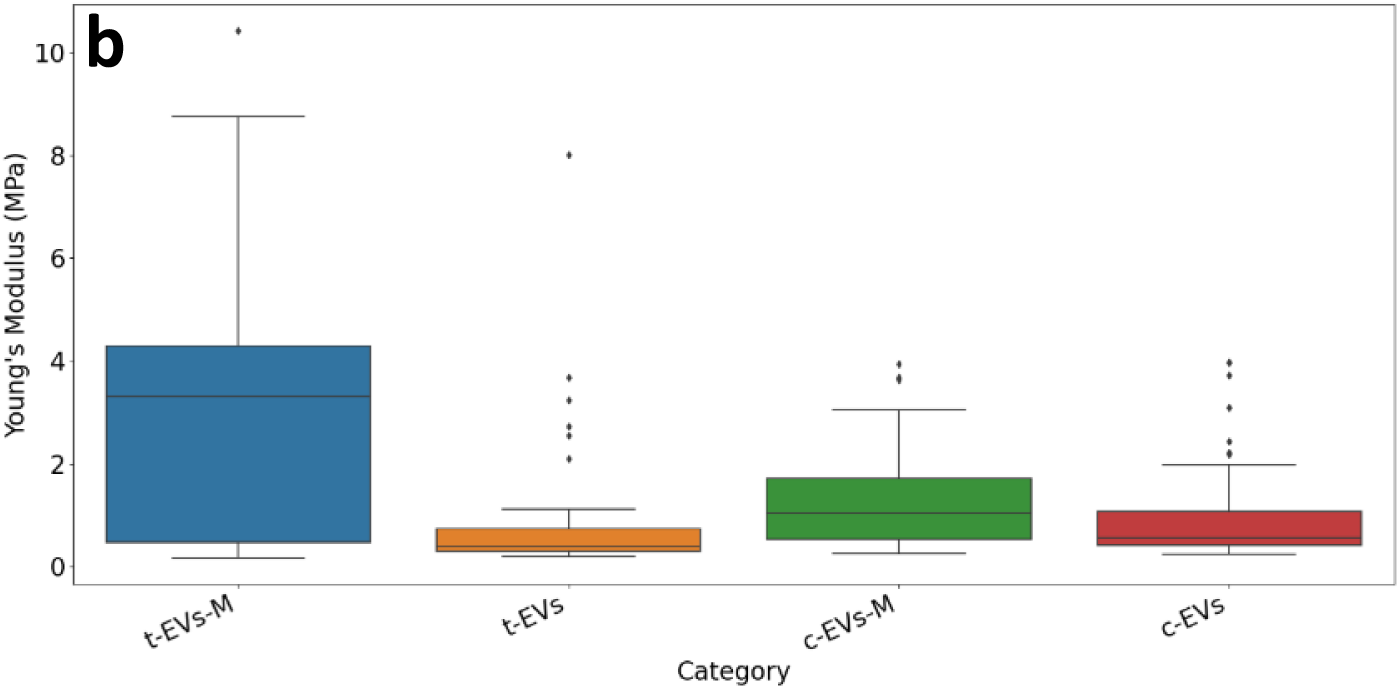
Young’s Modulus distribution of EVs derived from cells, in both conditions (control and treated samples). a. 3D plot of categorized distribution of Young’s modulus values on single EVs. Here each dot represents one EV. b. Box plot of all the average Young’s modulus values obtained for different EV subpopulations. For this analysis, we measured 102 EVs coming from the treated condition and 114 EVs coming from the control condition. Blue: t-EVs-M; Orange: t-EVs; Green: c-EVs-M and Red: c-EVs.

In Figure 7b, the box plot shows the overall distribution of average Young’s modulus values for the different EV subpopulations. t-EVs-M exhibited a broader range of Young’s modulus values compared to the other subpopulations, indicating increased mechanical heterogeneity. While mitochondria-containing lEVs tended to display higher elastic modulus than those lacking mitochondria, this difference cannot be attributed solely to the presence of mitochondria. It should also be noted that MitoTracker Green (MTG) can label not only intact mitochondria but also mitochondrial fragments. Other factors, such as variations in lipid composition or cargo organization, or multilayered structures (supplementary figure S3, showing images obtained by cryoEM), may also contribute to the observed mechanical differences. This hypothesis is supported by our previously published Raman spectroscopy results, which revealed notable changes in the lipid spectral signatures of EVs derived from B[a]P-treated cells, consistent with stress-induced membrane remodeling [27]. Quantitatively, the mean Young’s modulus of mitochondria-containing EVs was 3.09 MPa for t-EVs-M and 1.25 MPa for c-EVs-M, whereas EVs lacking mitochondria were overall softer, with mean values of 1.03 MPa and 0.96 MPa for t-EVs and c-EVs, respectively.

The resulting Young’s modulus distribution profile helped in observing the mechanical heterogeneity within the population of EVs. Bairamukov et al. also observed this heterogeneity within individual vesicles and in the lEVs subpopulation derived from blood plasma. Using AFM, they measured a Young’s modulus of 22.48 MPa [10]. In another study, the Young’s modulus of the honey derived EVs (sEVs) was determined to be in between ∼3– 165 MPa [11], this distribution suggests the presence of several EVs subpopulation present in the sample. Various cell sources and their physiological states affect the mechanical characteristics of extracellular vesicles. As was observed by Feng et al., they studied the nanomechanical signature of the EVs originating from blood and bone marrow of healthy donors and different cancer type patients. Not only did they measure the difference between the EVs coming from different sources from one patient, but also different Young’s modulus was observed from healthy donor and cancer patients (myeloma and lymphoma) [13]. These findings underscore the relevance of nanomechanical measurements of EVs for identifying biophysical alterations associated with disease and stress responses.

An important aspect when calculating the Young’s modulus values is to consider the indentation model applied. Normally the indentation model is chosen according to the shape of the AFM tip utilized and the adhesion forces between tip and the samples. In this study, as in the works of Bairamukov et al. and Karabasz et al., we used DMT model to qualify mechanically EVs subsets [10], [35]. DMT indentation model is based on hertz theory to calculate the Young’s modulus, this model was suitable for spherical shaped AFM tip used where the adhesion force between the tip and the sample was very small [1]. To estimate the elastic modulus (Young’s modulus), the linear segment of a retract curve recorded during scanning was fit automatically using the DMT model. This approach assumes a homogeneous mechanical response across the vesicle surface [36], which—as revealed by our findings—is an idealization that overlooks the intrinsic heterogeneity of EVs. Indeed, previously published articles have shown that EVs, due to their own structural composition, have intrinsically both linear and non-linear contact mechanics [26], [37], [38], [39], [40]. However, in our experiments, the indentation depth remained within ∼10% of the vesicle height, ensuring that the mechanical response stayed within the linear elastic regime, where the DMT model remains valid. Other models like thin shell and Canham Helfrich theory based models have been used for EVs; in these models the mechanical properties are dependent on the size of the EVs [26], [37], [38]. Indeed, Hertz contact, thin shell, modified Canham-Helfrich models and DMT, represent models that have been subject to “adaptations”/“corrections” in the study of extracellular vesicles [36]. However, it was noted that none of these models consider the inherent inhomogeneity of extracellular vesicles and that other and new more suitable models would be needed.

Nevertheless, despite their limitations, these mathematical models consistently offer the potential discrimination based on biomechanical diversity within EV populations and other kind of nanoparticles [36]. In recent articles, novel approaches such as nanomechanical means for discriminating vesicle subtypes, underline the great dynamics of this field of research. Piontek & Roos proposed a nanoindentation approach [41], [42] that was applied to extracellular vesicles by Vorselen et al., based on Canham-Helfrich theory, to discriminate lipoproteins from other nanoparticles, for example during extracellular vesicles isolation procedures[26]. Ridolfi et al, published their approach that they call “a crude but effective” one, to “leverage nanomechanics to discriminate” EV and lipid nanoparticles “that cannot be resolved by size alone”. They called their method “a morphometry assay” for the “rapid nanomechanical screening of mixed nanoparticles samples” [9]. Our work follows the same rationale—applying a “crude but efficient” nanomechanical strategy, with its inherent compromises, to discriminate between nanovesicles of similar dimensions but differing in internal composition and mechanical behaviour.

The AFM study of soft nanoscaled objects in liquid requires reaching a compromise between applied force, tapping parameters, AFM probe geometry and reliable morphology. Each model is based on the assumptions that might not necessarily be applicable for the EVs, but all the models are able to reveal the heterogeneity among the EV populations. Ultimately, our objective is not to assign absolute mechanical values to specific EV subpopulations but to demonstrate that nanomechanical profiling can serve as a robust and label-free tool for distinguishing, screening, and classifying EV subsets within complex biological samples.

Another crucial aspect of this approach lies in data analysis. In recent years, statistical and machine-learning methods have become prevalent for accelerating and improving the accuracy of data interpretation [43]. These methods have also shown the potential to identify the correlations within large datasets and identify the crucial features in the data [44]. The machine learning algorithms have already been used to classify cell surfaces of biological cells [45] and there is a great potential to use the same approach to classify and characterize EV subpopulations [46].

### 3. Statistical analysis

LDA was applied to visualize the discrimination between different EV subpopulation where we used Young’s modulus distribution, height, and the maximum diameter as the features for each EV. LD1 and LD2 are linear discriminants derived from the LDA transformation, these discriminants helped in the maximizing the variance between different population while reducing the variance within each population.

In Figure 8, the first curve is the distribution of each population according to LD1. Here, we can clearly observe the separation of EVs with mitochondria from EVs without mitochondria. Hence, we decided to perform ANOVA on LD1 scores of the EV with mitochondria (t-EVs-M and c-EVs-M) and EV without mitochondria (tEVs and cEVs) with the assumption that with LD1 scores, we can identify the EVs containing mitochondria. We obtained p-value 9.47 10, indicating that the difference in LD1 is significant between the combined population of EVs containing or not mitochondria. Another hypothesis tested with ANOVA was that control and treated conditions could be discriminated for EVs containing mitochondria. We used LD2 scores of t-EVs-M and c-EVs-M, we obtained a p-value 1.1 10. These results confirm that with LDA on three variables - Young modulus values, maximum diameter and height of lEVs - we can differentiate between the lEVs populations while ANOVA further confirms the significance of the differences observed.

**Figure 8.**
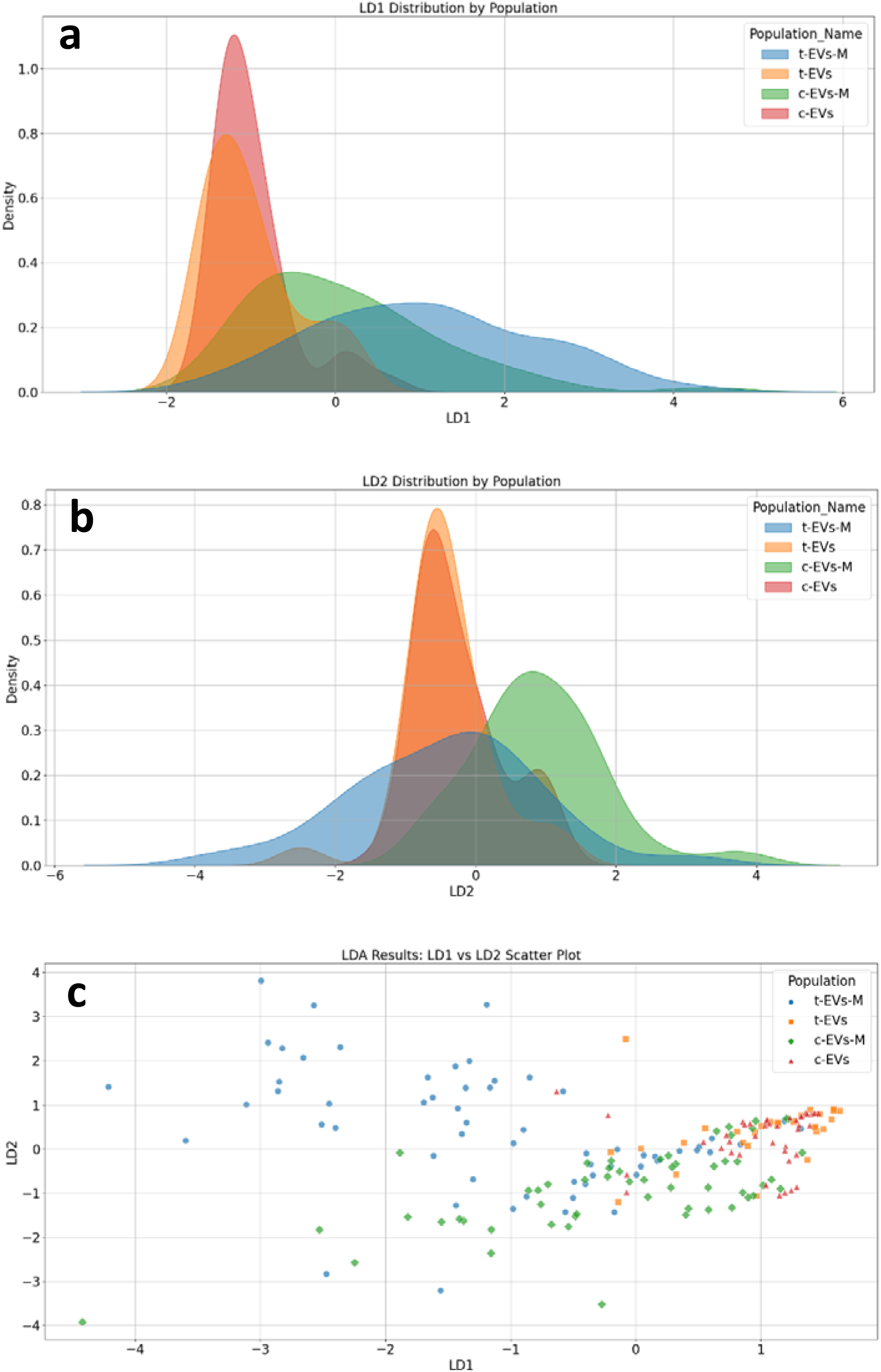
LDA analysis for EV discrimination. a and b are distribution curves based on LD1 and LD2 respectively. c is the scatter plot between LD1 and LD2 score where each point in the plot represents an individual EV. Blue: t-EVs-M; Orange: t-EVs; Green: c-EVs-M and Red: c-EVs.

After confirming the reliability of morpho-mechanical data (Young’s modulus, height, and the maximum diameter) to discriminate different EV subpopulations, we applied the same assumptions to the Random Forest algorithm. This algorithm creates multiple decision trees using random subsets of data and features, combining results to improve accuracy and reduce overfitting. We tested this model using LOOCV where n-1 data points are used for training, and the remaining single data point is used for testing, repeating this process for all data points. The confusion matrices (Figure 9) present the results obtained using this classification model. The model achieved an accuracy of 84.21% in identifying EVs containing mitochondria from others (Assumption 1). Additionally, it achieved 76.19% accuracy in distinguishing c-EVs-M from t-EVs-M (Assumption 2).

**Figure 9.**
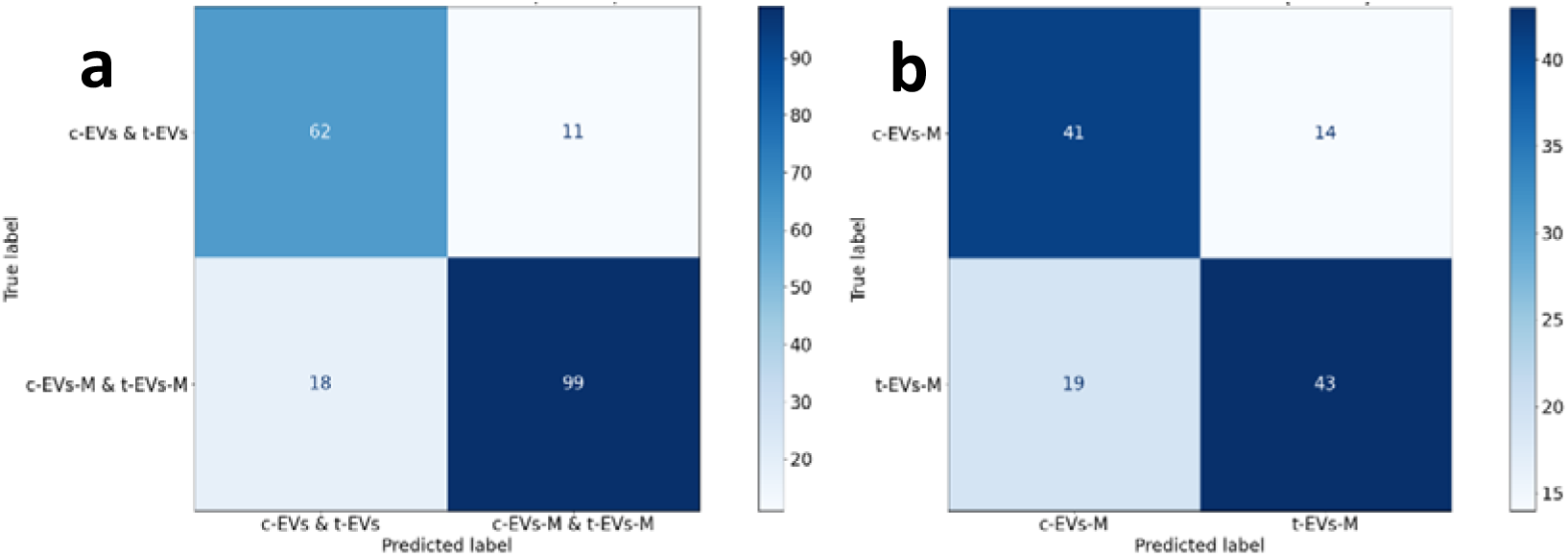
Confusion matrices of the classification of EV subpopulations, based on lEVs young modulus values, maximum diameter and height. a. Classification results for identifying EVs containing mitochondria. b. Classification results for distinguishing between c-EVs-M and t-EVs-M.

### Conclusion

This study explored a way to discriminate EVs originating from HMEC-cells exposed to B[a]P or not through morpho mechanical properties. The discrimination was based on the measurements obtained at single vesicle level. The elastic modulus of EVs offered insights into both their physiological state and the culture condition of the parent cells. By examining the mechanical properties of the EVs, we could determine their membranous nanomechanical properties and differentiate distinct EV subpopulations.

Using a combination of fluorescence microscopy and atomic force microscopy, we were able to measure accurately the mechanical properties of EVs containing mitochondria or not. Our work shows that by adopting this low-stringency approach (random points), it is possible to discriminate between vesicles containing mitochondria and those not, and even between vesicles containing mitochondria but derived from control or B(a)P-treated cells. Indeed, our findings revealed that EVs containing mitochondria from B[a]P-treated cells exhibited higher apparent elastic modulus values (3.09 MPa) compared to those from control cells (1.03 MPa).

Through statistical methods such as LDA and ANOVA, we demonstrated the ability to discriminate between EVs containing mitochondria and those without, showing a significant difference in morpho-mechanical properties (p-value = 9.5 × 10⁻²⁰). Additionally, we confirmed the possibility of distinguishing the source of mitochondria-containing EVs (control vs. treated) with a p-value = 1.1 × 10⁻⁸. The results from the Random Forest model further validate the effectiveness of this approach in accurately identifying and classifying EV subpopulations. The high accuracy achieved emphasizes the potential of morpho-mechanical data for robust classification, particularly in identifying mitochondria-containing EVs and differentiating their sources.

There remains an ongoing debate in the literature regarding the optimal balance between applied force, scanning parameters, probe geometry, morphology preservation, and data-processing models in AFM-based nanomechanics. Our work fits within this framework, supporting the concept that nanomechanical profiling represents a powerful analytical approach to discriminate nanovesicle subpopulations of identical size within complex biological samples. Our work also shows that fluorescence would no longer be necessary to distinguish lEVs containing mitochondria and coming from treated cells.

Future improvements as immuno-enrichment on the chip can further aid in selecting specific EV subpopulations. We emphasize that such study could be envisioned as a nanoanalytical method to assess the impact of various toxicological pollutants on EVs subsets and may also serve as a diagnostic tool for a wide range of medical applications.

## Acknowledgments and fundings

This work has been funded by the EIPHI Graduate School (contract ANR-17-EURE-0002) and Bourgogne-Franche-Comté Region. This work was supported by the French National Agency for Research (ANR; ENDOMITOPAH project: R23137NN). The authors wish to thank the Biophysics and Structural Biology Core Facility of Service Mutualisé de Plateformes (SMP) - Pôle BMS. (Université de Lorraine-http://umsibslor.univ-lorraine.fr), the laboratory staff of the Clinical-Innovation Proteomic Platform (CLIPP) and Nano2BIO team at FEMTO-ST Institute for fruitful discussion and technical/instrumental supports.

