## Supplementary Figures for "Large extracellular vesicles subsets and contents discrimination: the potential of morpho mechanical approaches at single vesicle level"

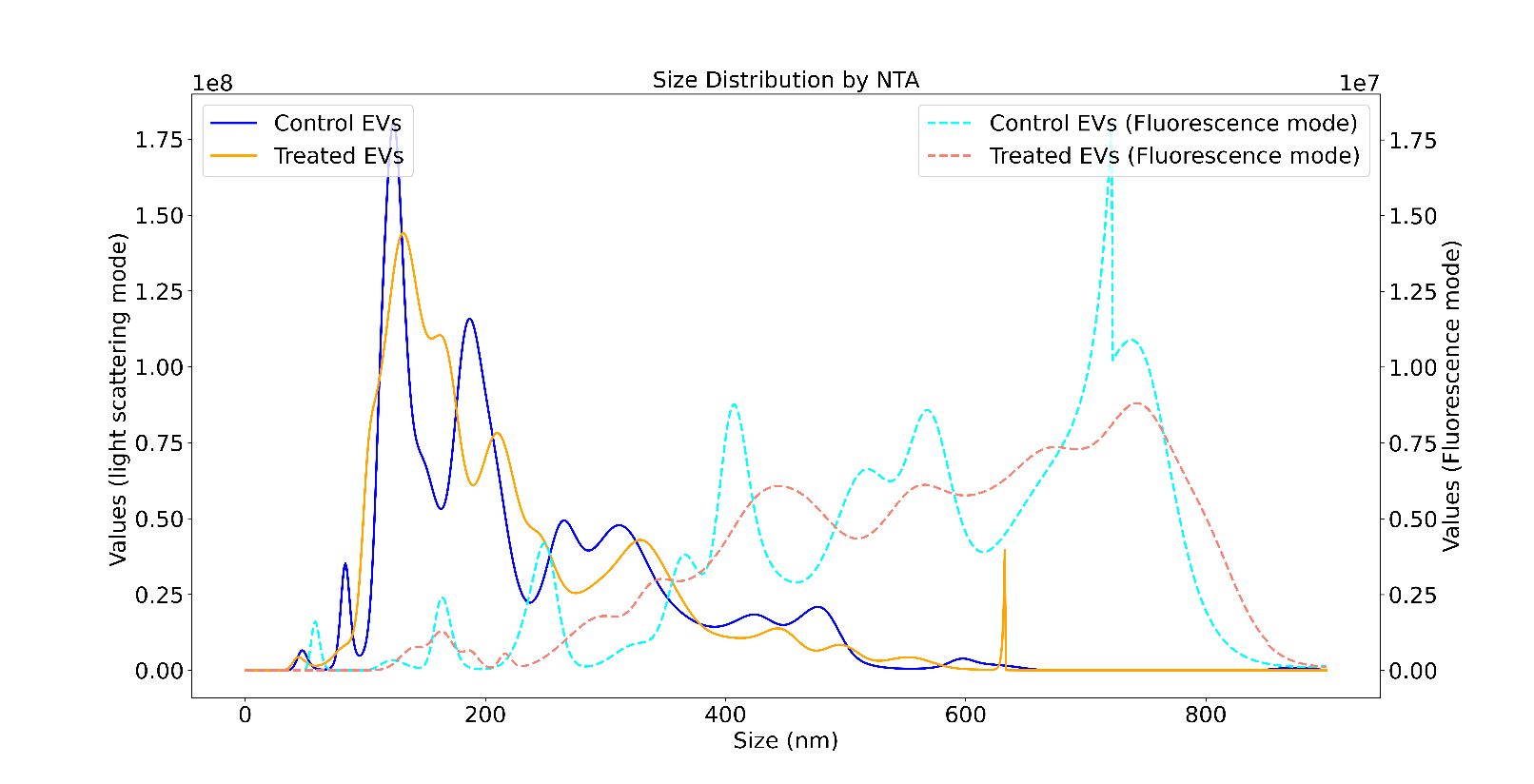

**Figure S1.** NTA results showing the concentration of particles as a function of their size for control and treated samples.

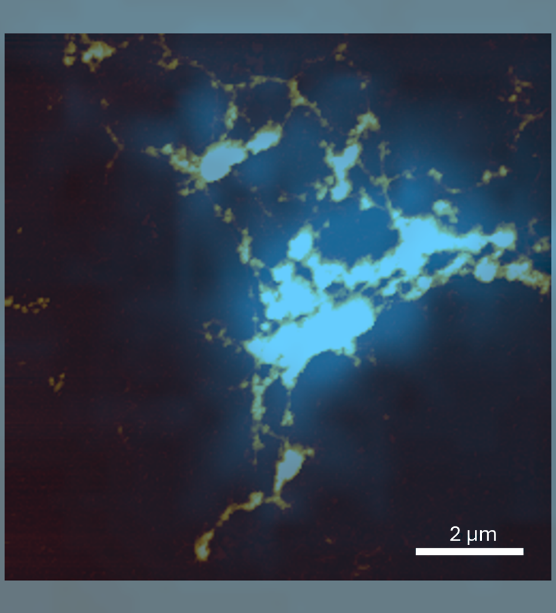

**Figure S2.** Overlaid fluorescence and AFM image (figure 4e). This overlaid image shows the presence of mitochondria inside these vesicles.

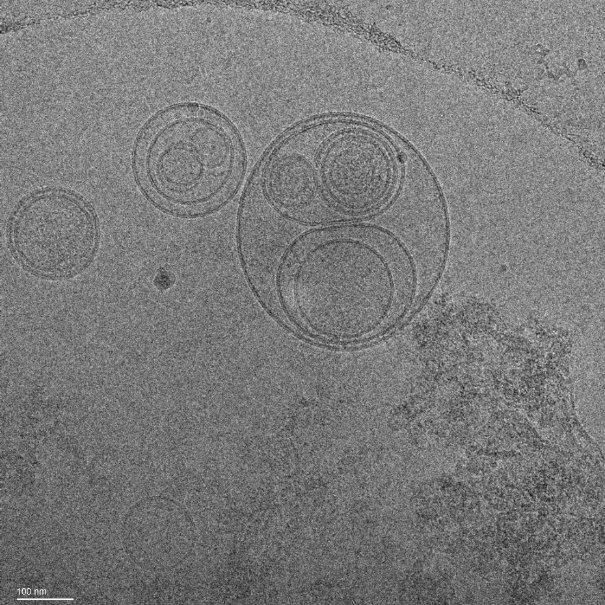

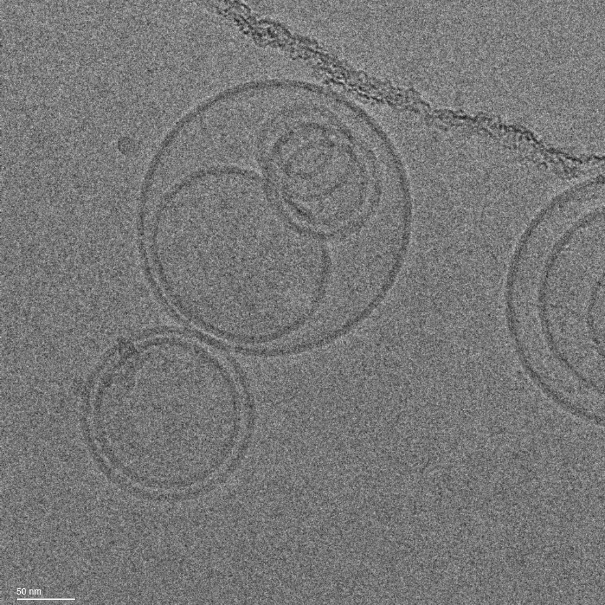

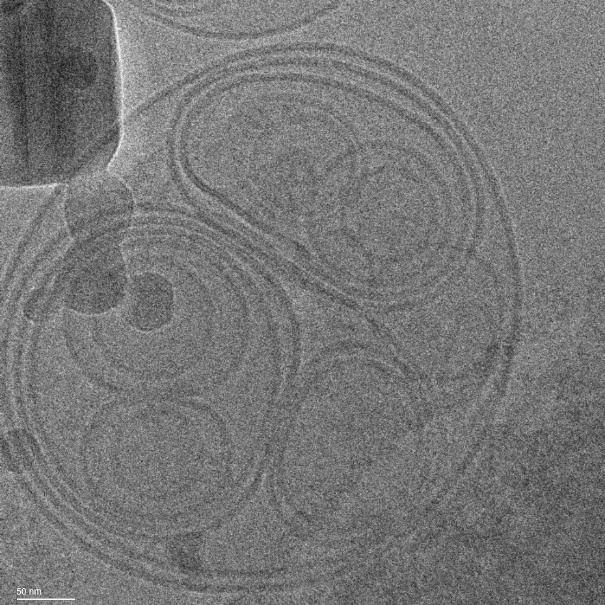

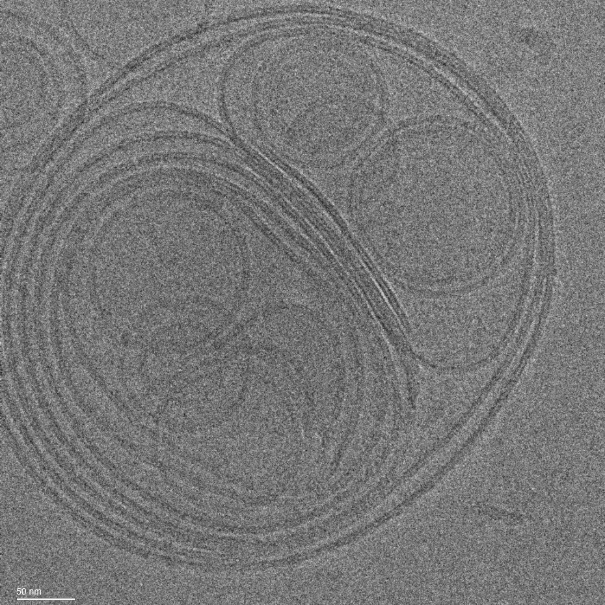

**d**

**c**

**a**

**b**

**Figure S3.** Cryo-EM images showing different EV subpopulations from treated condition, including single bilayer vesicles and multilamellar (multiple-layer) vesicles. Scale bar = 100 (in a) and 50 nm (b, c, d).
